# Transcranial focused ultrasound modulates visual thalamus in a nonhuman primate model

**DOI:** 10.1101/2024.08.05.606669

**Authors:** Kai Yu, Bin He

**Affiliations:** Department of Biomedical Engineering, Carnegie Mellon University, Pittsburgh, PA 15213; Neuroscience Institute, Carnegie Mellon University, Pittsburgh, PA 15213

**Keywords:** Transcranial focused ultrasound neuromodulation, nonhuman primate, thalamus, extrastriate visual cortex, electroencephalography, source imaging, local field potential

## Abstract

The thalamus plays a pivotal role as a neural hub, integrating and distributing visual information to cortical regions responsible for visual processing. Transcranial focused ultrasound (tFUS) has emerged as a promising non-invasive brain stimulation technology, enabling modulation of neural circuits with high spatial precision. This study investigates the tFUS neuromodulation at visual thalamus and characterizes the resultant effects on interconnected visual areas in a nonhuman primate model. Experiments were conducted on a rhesus macaque trained in a visual fixation task, combining tFUS stimulation with simultaneous scalp electroencephalography (EEG) and intracranial recordings from area V4, a region closely linked to the thalamus. Ultrasound was delivered through a 128-element random array ultrasound transducer operating at 700 kHz, with the focus steered onto the pulvinar of the thalamus based on neuroanatomical atlas and individual brain model. EEG source imaging revealed localized tFUS-induced activities in the thalamus, midbrain, and visual cortical regions. Critically, tFUS stimulation of the pulvinar can elicit robust neural responses in V4 without visual input, manifested as significant modulations in local field potentials, elevated alpha and gamma power, corroborating the functional thalamocortical connectivity. Furthermore, the tFUS neuromodulatory effects on visually-evoked V4 activities were region-specific within the thalamus and dependent on ultrasound pulse repetition frequency. This work provides direct electrophysiological evidence demonstrating the capability of tFUS in modulating the visual thalamus and its functional impact on interconnected cortical regions in a large mammalian model, paving the way for potential investigations for tFUS treating visual, sensory, and cognitive impairments.

## Introduction

The thalamus is a critical neural hub, acting as a principal relay station that channels sensory data to various cortical areas responsible for processing visual cues^1,2^. This critical role of the thalamus to integrate and distribute visual information renders it an essential focus for studying neural mechanisms and interventions that could potentially enhance cognitive functions^3,4^ and treat neurological disorders^5^ and brain injury^6,7^.

Recent advances in neuromodulation technology have spotlighted transcranial focused ultrasound (tFUS) as a promising tool for non-invasive brain stimulation. tFUS provides a unique advantage by offering high spatial precision while targeting deep brain structures without necessitating surgical intervention^8^. This modality leverages focused ultrasound waves to modulate neuronal excitability, thereby influencing neural circuits in a controlled manner, impacting neural connections, and potentially altering behavioral outcomes. Although studies using rodent models have significantly contributed to our understanding of tFUS’s diverse parameters and their neuromodulatory effects^9–14^, translating these findings to human applications is challenging due to fundamental differences in anatomy between rodents and humans. Recent research involving nonhuman primates (NHP) has shown promising results, with tFUS successfully modifying behaviors like response times in antisaccade tasks^15^ and decision-making processes^16^ by targeting the frontal eye field. Furthermore, tFUS has proven effective in influencing processes such as credit assignment and value assessment within specific brain regions including area 47/12o and the anterior cingulate cortex^17^. It can also adjust localized brain activities, transform interactions within brain networks^18,19^, and even curb epileptiform activities and decrease the frequency of acute epileptic seizures^20^. These advancements highlight the tFUS to serve as an influential neuromodulatory approach in larger mammals.

Our recent investigations have successfully demonstrated that tFUS remotely modulates an extrastriate visual cortical region V4 by stimulating the frontal eye fields (FEF) through single neuronal activities and local field potentials^21^. These findings show that tFUS can affect brain regions that are not directly targeted by the ultrasound beam but are functionally interconnected.

Building on this foundational work, the current study seeks to explore and expand the application of tFUS to the visual thalamus in an NHP model trained to perform a visual task. The primary objective is to ascertain the feasibility of localizing tFUS-induced neural activity within the thalamus and to verify these effects using noninvasive neuroimaging and intracranial neural electrophysiological recording techniques. Specifically, we employ simultaneous 12-channel scalp electroencephalography (EEG) for source imaging and 96-channel intracranial recordings from the V4 area—a region closely linked with the thalamus—on a head-fixed, behaving rhesus macaque monkey to validate the observed EEG source imaging results.

## Material and methods

### Experimental Subject

We conducted recordings using an electrode array implanted in the visual cortex of a rhesus macaque (*Macaca mulatta*), simultaneously applying transcranial focused ultrasound stimulation (tFUS). The macaque was trained for a passive fixation task where it maintained gaze on a central point while a natural image appeared for 400 ms in the receptive field of area V4. Subsequent to this fixation, the animal executed a visually guided saccade to conclude the trial (Fig. 1a). In the neuromodulation session, we alternated between 120 trials with only visual stimulation and 120 trials with both visual and tFUS stimulations, all randomized (Fig. 1b). In the tFUS only sessions without actual visual stimuli presented on the screen after valid eye fixation, we randomly assigned the tFUS stimulation to half of the overall 240 trials. This research protocol received approval from the Institutional Animal Care and Use Committee at Carnegie Mellon University.

**Figure 1.**
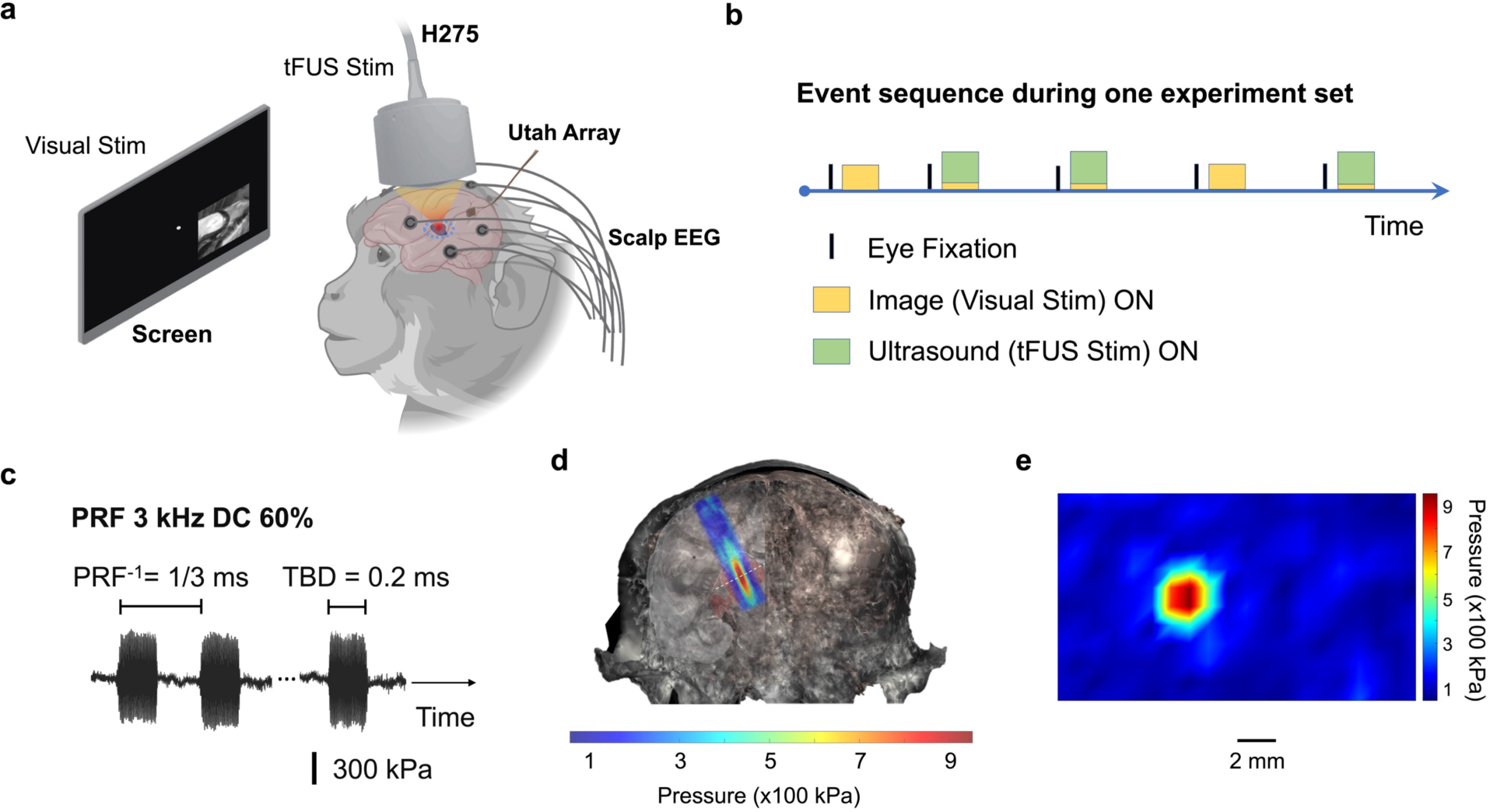
**a**, The experimental setup for concurrent scalp electroencephalography (EEG) recordings and intracranial neural recordings through a Utah array implanted in area V4 during visual and/or transcranial focused ultrasound (tFUS) stimulations. The ultrasound focus was steered axially and targeted at the thalamus (RAS coordinates: −6.56, −11.25, 3.83). **b**, Event sequence during one experiment set with tFUS randomly occurring during half of the trials. The trials with ultrasound stimuli only (sonication duration: 400 ms) or coupled with visual stimuli (duration: 400 ms) in some experiment sets were delivered to the monkey after successful eye fixation were detected. **c**, The transcranial ultrasound temporal profiles employed a pulse repetition frequency (PRF) of 3 kHz with the burst duty cycle (DC) of 60%. The tone burst duration (TBD) was set at 0.2 ms. **d**-**e**, The characterizations of the ultrasound pressure field generated by the 128-element random array ultrasound transducer H275 in the presence of a fully hydrated monkey skull sample are illustrated along the axial direction (**d**) and in the lateral direction (**e**) at the location marked with a white dashed line in (**d**).

### Ultrasound Setup

We employed a custom 128-element random array ultrasound transducer (H275) operating at a fundamental frequency of 700 kHz and transmitted ultrasound from a big acoustic aperture (diameter: 60 mm, F-number: 0.58), manufactured by Sonic Concepts, Inc. (Bothell, Washington, USA). A holder for the transducer, featuring X and Y axis indicators and constructed from Tough 1500 Resin, along with a specialized ultrasound collimator made from Clear Resin, were both 3D printed using a Form 3+ printer (Formlabs, Somerville, MA, USA). These components secured the transducer in a fixed position on the animal’s chair and aided in directing the ultrasound beam along precise trajectories. The H275 transducer was driven by a Vantage 256 research ultrasound system (Verasonics, Kirkland, WA, USA) to generate specific ultrasound pulse repetition frequency (PRF, 3 kHz or 40 Hz) and burst duty cycle (DC, 60% or 8%) (Fig. 1c).

We generated a full skull model of the NHP from CT scans. By aligning the MRI-based brain model of the subject with the skull model, we adjusted the transducer’s position on the scalp to accurately direct the tFUS beam to a thalamic area, pulvinar, using the Macaque NeuroMaps Atlas^22^ (Fig. 1). This alignment provided spatial guidance to improve the precision of tFUS targeting during live experiments. To evaluate the ultrasound transmission performance and analyze temporal and spatial dynamics near the brain targets, we created a three-dimensional *ex vivo* pressure mapping system. This system featured a needle hydrophone (HNR500, Onda Corporation, Sunnyvale, CA, USA) submerged in degassed water and driven by a 3-axis positioning stage (Velmex, Inc., Bloomfield, NY, USA) to map the spatial-temporal pressure/intensity field of ultrasound after passing through a piece of fully hydrated *ex vivo* monkey skull bone. The needle hydrophone was positioned beneath the skull sample, measuring ultrasound pressure at discrete locations with a scanning resolution of 0.5 mm. The transcranial spatial peak pressure magnitude was measured as 950.9 kPa (spatial-peak pulse-average intensity I_SPPA_: 6.99 W/cm^2^ when the ultrasound pulse duration is 200 μs) with an axial focal length of 13.9 mm (−6dB contour, Fig. 1d) and a lateral focal width of 2.4 mm (−6dB contour, Fig. 1e). For precise focus steering at the cortical or deep brain, we aligned the X-axis of the H275 with the brain’s medial-lateral axis.

### Electrophysiological Recordings and Preprocessing

To monitor the whole brain responses to the tFUS stimulation at visual thalamus, 12 gold-cup scalp electroencephalography (EEG) electrodes were placed over the shaved monkey head (Fig. 1a), and their spatial locations were digitized with a 3D digitizer (FASTRAK, Polhemus, Colchester, VT) to inform further EEG source imaging (ESI)^23^. To further examine the local brain activations due to the thalamus stimulation, we harnessed the neural connection between the thalamus and an extrastriate visual cortex V4, and recorded neural activities through a 96-channel “Utah” electrode array (Blackrock Neurotech, Salt Lake City, UT, USA) implanted in the V4 area on the prelunate gyrus, situated medial to the inferior occipital sulcus (see Stan et al. for more details of the array implantation in this animal RA^24^).

The scalp EEG and intracranial electrophysiological signals were acquired by a Ripple recording system and Trellis software suite at a sampling frequency of 30 kHz (Ripple Neuromed, Salt Lake City, UT, USA) with a bandpass filter of 0.3 Hz to 7.5 kHz set by the hardware for raw data acquisition, and additionally band passed with a range of 0.3 to 250 Hz for saving both EEG and local field potentials. The EEG channels were further band-pass filtered with a lower cut-off frequency at 1 Hz and the higher cut-off frequency at 50 Hz. The local field potential (LFP) data were further high-pass filtered during postprocessing in the FieldTrip toolbox^25^ with a cut-off frequency of 1 Hz and notch filtered at 60 and 120 Hz to remove powerline noise.

### Neural Electrophysiological Data Analysis

After the neural datasets were preprocessed, they were further segmented into epochs based on the visual/ultrasound stimulation events. For EEG data, the pre-stimulus period was set as 200 ms before an event trigger signal, and the 500 ms period after the trigger signal was deemed as post-stimulus period in EEG individual epoch. The 0.7-s EEG epochs were then averaged across the trials with the same experimental condition by aligning specific trigger events. For EEG source modeling and imaging, the boundary element method (BEM) head model^26,27^ for the monkey was established using CURRY (Version 8.0.6, Compumedics USA, Charlotte, NC, USA), which consisted of three layers, i.e., scalp, skull and brain with relative conductivities of 0.33, 0.0125 and 0.33, respectively. The minimum norm (MN) and standardized low-resolution brain electromagnetic tomography (sLORETA)^28^ were used to solve the inverse problem and reconstruct the source activities from cortical and deep brain regions. For visualization purpose, 75% thresholding was applied in the source images.

To compare LFP temporal waveforms across different conditions within the time window from - 0.5 to 1 second, we averaged the epoch data across all trials at representative channels for each experimental set, while retaining individual trial data. This approach enabled nonparametric statistical analyses using a temporal cluster-based permutation test, with Monte-Carlo estimates of significance probabilities derived from the permutation distribution via the *ft_timelockstatistic* function in the FieldTrip toolbox^25,29^. Color-coded, vertical shaded bars indicated time segments with significant differences between two specified experimental conditions. The shaded areas behind the mean LFP profiles represent ± one standard error of the mean. To visualize LFP changes, we generated topographies of t-statistics at specific time frames using the *ft_timelockstatistic* process, highlighting clusters across all 96 channels of the Utah array, with a significance level set at 0.05. Gray asterisks marked electrode clusters with significant differences in the electrode array topographic maps. We used 5000 draws from the permutation distribution for all tests, which were two-sided to identify both positive and negative clusters in the temporal and spatial domains. The time-frequency analyses producing the power-spectra were also implemented in the FieldTrip toolbox via the *ft_freqanalysis* function adopting multi-taper time-frequency transformation based on multiplication in the frequency domain with frequency step size of 0.1 Hz, time step size of 1 ms, and spectral smoothing through multi-tapering.

## Results

### EEG source localization of tFUS induced brain activity at frontal eye field

To test the feasibility of using scalp EEG source imaging for mapping and localizing the *in vivo* brain responses to tFUS stimulation, we recorded the brain activations in response to the excitatory tFUS stimulation (PRF: 3 kHz, burst duty cycle: 60%) targeted at the FEF of a monkey brain without the concurrent presentation of visual stimuli following the protocol described in our previous work on the monkey subject^21^. Fig. 2 depicts the findings from the EEG recordings and cortical brain source analyses. Fig. 2a presents a butterfly plot of the 12-channel scalp EEG signals averaged from 120 trials only with tFUS stimulations (sonication duration: 400 ms). Each color represents an individual EEG channel, displaying the temporal dynamics of the EEG responses to the ultrasound stimulation. The selected two time points of 48 and 75 ms represent the local peaks in the mean global field potential of the EEG (dashed lines). The EEG-based source localization off brain activity in response to tFUS stimulation was performed using the subject-specific 3D head model at those two specific time points after the onset of tFUS stimulation (Fig. 2b and 2d). Voltage topographic maps with contour lines illustrate the positions of the EEG electrodes over the scalp at these time points.

**Figure 2.**
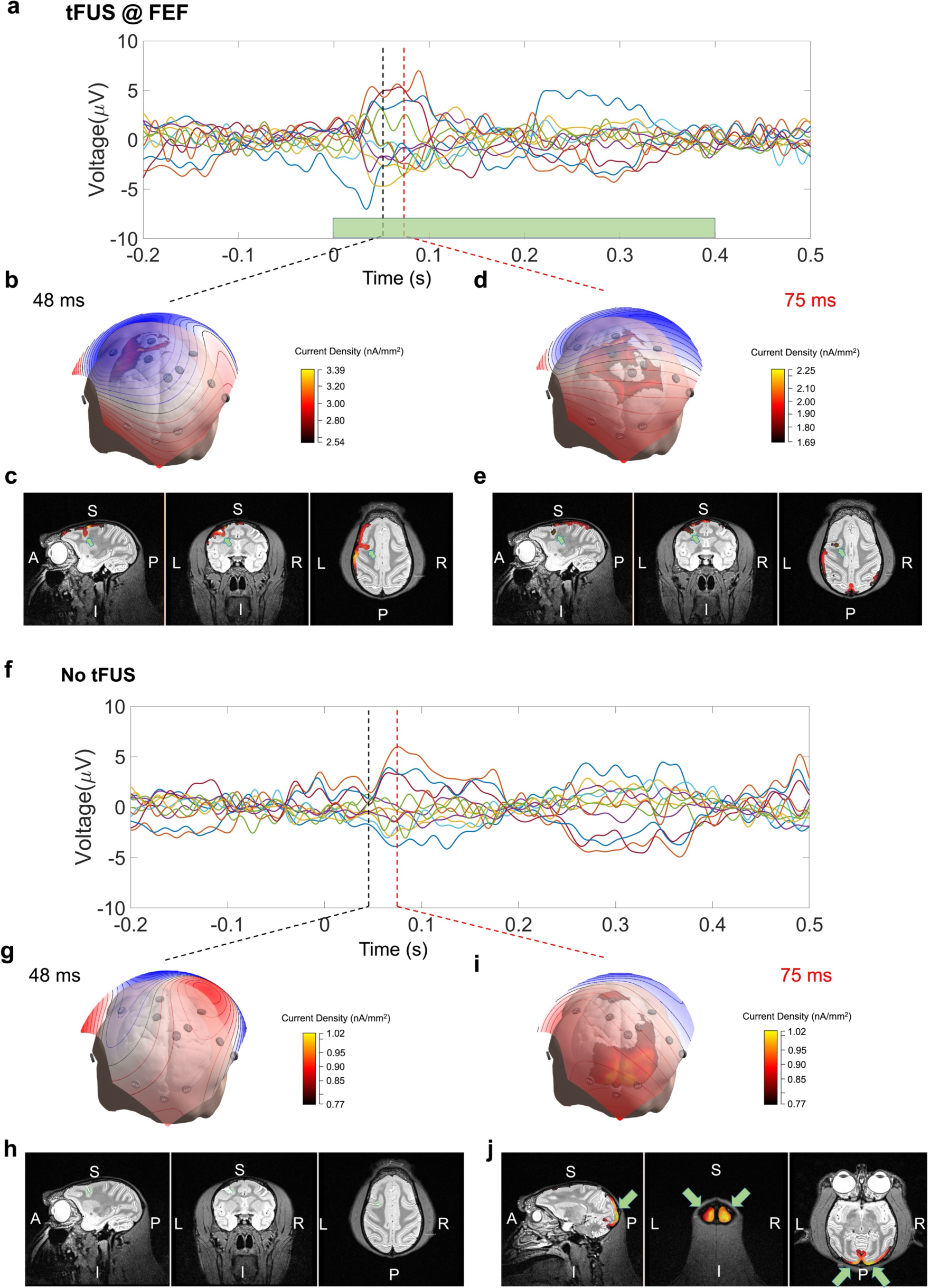
Scalp EEG monitoring the brain activations in response to tFUS stimuli delivered to frontal eye field (FEF) of the monkey brain without coupling with visual stimuli presented on the screen. **a**, A butterfly plot illustrating the 12-channel scalp EEG signals averaged from 120 trials with only ultrasound stimulations (sonication duration: 400 ms). Each color of the line profiles represents one EEG channel. **b**-**e**, The EEG source localization in responding to the FEF stimulation with tFUS in a 3D head model (**b, d**) at two time points of 48 and 75 ms. The locations of EEG electrodes over the scalp and the voltage topographic maps with contour lines at those two time points are illustrated over the surface of the 3D head model. The source images acquired from minimum norm estimations along the sagittal, coronal and horizontal planes of the brain magnetic resonance images further delineate the localizations of tFUS-evoked source activities at the cortical brain. The source activities reconstructed at prearcuate sulcus are observed (**c**, **e**) (indicated with green arrows), where the FEF is located. **f**, Butterfly plot illustrating the 12-channel scalp EEG signals averaged from 120 trials without tFUS or actual visual stimulus presented to the monkey subject. Individual EEG channels are illustrated with different line colors. **g**-**j**, The EEG source localization without presenting external stimuli at the same two time points of 48 (**g**) and 75 ms (**i**). Similarly, the source estimations on top of the brain magnetic resonance images along the sagittal, coronal and horizontal planes indicate absences of observable cortical activities at 48 ms (**h**), while appearance of strong visual cortical activities at 75 ms (**j**) (indicated with green arrows) potentially due to attention and/or visual anticipation in this trained visual paradigm.

Further, source images acquired from minimum norm estimations along the sagittal, coronal, and horizontal planes of the brain magnetic resonance images delineate the localizations of tFUS-evoked source activities at the cortical level. The source activities reconstructed at the prearcuate sulcus, where the FEF is located, are observed with current source density (CSD) ranging from 2.9 - 3 nA/mm^2^ at 48 ms (Fig. 2c) and reduced to 1.69 - 1.80 nA/mm^2^ at 75 ms (Fig. 2e, indicated by green arrows).

To establish a baseline thus a comparison, we recorded EEG signals from 120 trials without tFUS or any visual stimuli, but only valid eye fixations detected. The butterfly plot in Fig. 2f illustrates these baseline EEG signals across those 12 channels. Fig. 2g and 2i display the EEG source localization results at 48 ms and 75 ms, respectively. The source estimations overlaid on the brain magnetic resonance images along the sagittal, coronal, and horizontal planes show an absence of observable cortical activities at 48 ms (Fig. 2h). However, at 75 ms, localized visual cortical activities are noted with a maximum CSD of 1.02 nA/mm^2^ presented at the primary visual cortex, and a range of CSD from 0.77 to 0.85 nA/mm^2^ localized to extrastriate corticies, which might be due to attention^30^ and/or visual expectation^31^ in this trained visual paradigm (Fig. 2j, indicated by green arrows).

Notably, a late EEG signal complex starting after 200 ms post the onset of stimulation or expected event can both be seen in both FEF stimulation (Fig. 2a) and no stimulation (Fig. 2f) conditions. The tFUS stimulation at the FEF is seen to shift this signal complex to be earlier for 35 ms than it is in the baseline condition, while the stimulation does not significantly alter the duration (145 ms) of this signal complex.

### EEG source localization of tFUS induced brain activity at thalamus

We further adjusted the experimental setup and steered the ultrasound focus onto thalamus using the aforementioned RAS coordinates of a Macaque brain atlas. The experiment was also conducted without visual stimuli presented on the screen. Fig. 3a presents a butterfly plot of the 12-channel scalp EEG signals averaged from 120 trials with only tFUS stimulations (sonication duration: 400 ms). Each line color represents a different EEG channel, displaying the temporal dynamics of EEG responses to the ultrasound stimulation.

**Figure 3.**
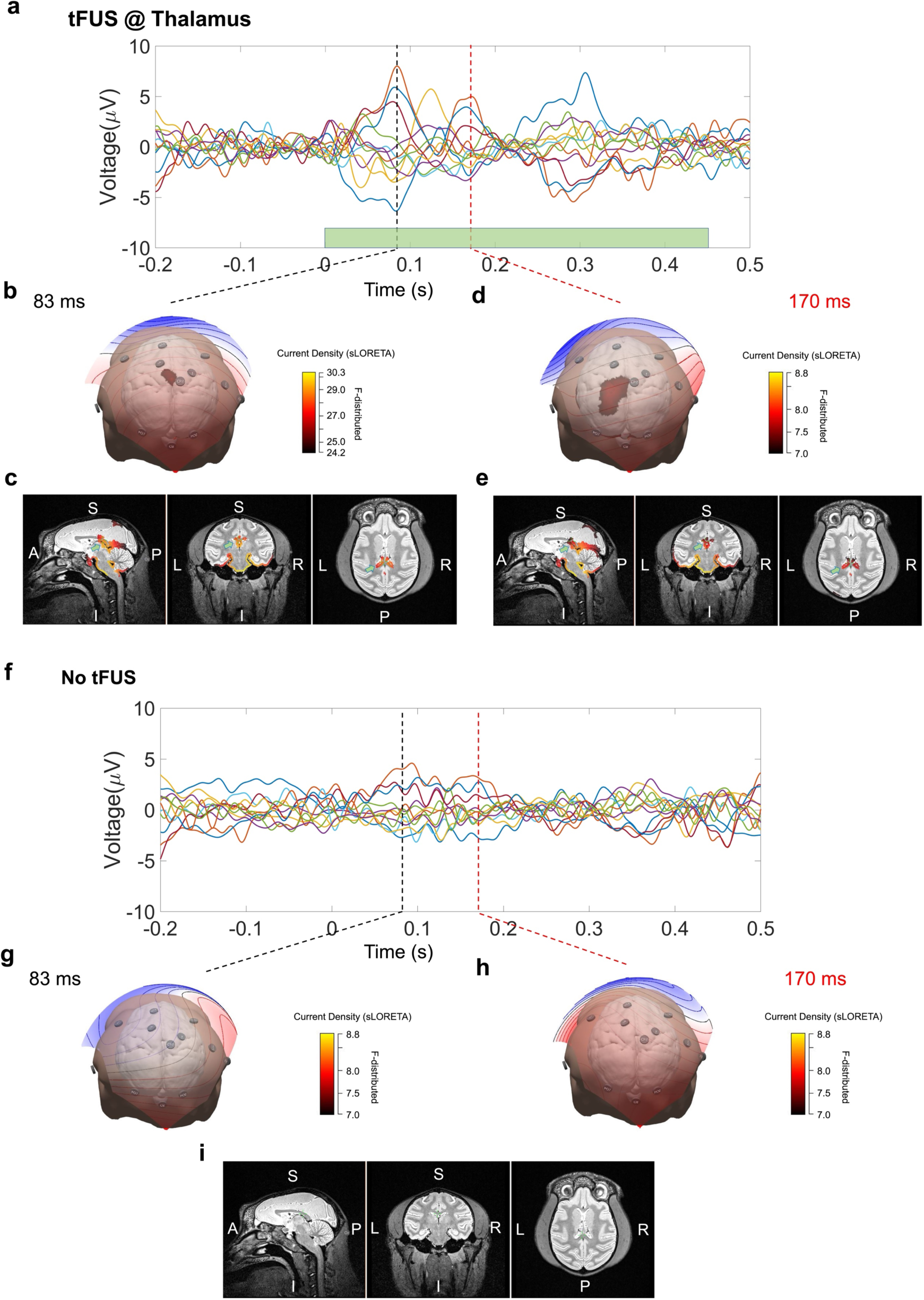
Scalp EEG monitoring the brain activations in response to tFUS stimuli delivered to the thalamus of the monkey brain without visual stimuli presented on the screen. **a**, A butterfly plot illustrating the 12-channel scalp EEG signals averaged from 120 trials with only ultrasound stimulations (sonication duration: 400 ms). Each color of the line profiles represents one EEG channel. **b**-**e**, The EEG source localization in responding to the thalamic stimulation with tFUS in a 3D head model (**b**, **d**) at two time points of 83 and 170 ms. The locations of EEG electrodes over the scalp and the voltage topographic maps with contour lines at those two time points are illustrated over the surface of the 3D head model. The source images reconstructed with standardized low-resolution electromagnetic tomography (sLORETA) method along the sagittal, coronal and horizontal planes of the brain magnetic resonance images further delineate the localizations of tFUS-evoked source activities at the cortical brain. Strong source activities at the thalamus are observed (**c**, **e**) (indicated with green arrows). **f**, Butterfly plot illustrating the 12-channel scalp EEG signals averaged from 120 trials without tFUS or actual visual stimulus presented to the monkey subject. Individual EEG channels are illustrated with different line colors. **g**-**i**, The EEG source localization without presenting external stimuli at the same two time points of 83 (**g**) and 170 ms (**h**). The source estimations on top of the brain magnetic resonance images along the sagittal, coronal and horizontal planes do not exhibit observable source activities at cortical or deep brain regions using the same color scale in the panel (**d**).

The EEG source localization in response to tFUS stimulation was performed using the same 3D head model at two specific time points, 83 ms and 170 ms post-stimulation (Fig. 3b and 3d). For an improved localization at deep brain structures, source images were reconstructed with the standardized low-resolution electromagnetic tomography (sLORETA) method^28,32^ at the sagittal, coronal, and horizontal planes of the brain magnetic resonance images. Specifically, at 83 ms (Fig. 3b-c), significant source activities at the thalamus are observed (F values: 28 – 30.3, indicated with green arrows), and some activations in adjacent midbrain areas, potentially involving regions such as the superior colliculus are also seen (F values: 29.2 – 29.7). There are areas of activation extending into the cerebellum (F values: 24.2 – 29.1), which may be involved due to its role in coordinating sensory information and motor control. There is also visible weak activation in the cortical regions (F values: 24.2 – 26.0), potentially involving parts of the parietal lobes. These areas could be engaged due to their connectivity with the thalamus and involvement in higher-order processing and attention. Furthermore, activation extends into the medial parts of the temporal lobes, which may include the hippocampus and other medial temporal structures that are involved in memory and spatial navigation. At a later time of 170 ms (Fig. 3d-e), besides the observed thalamic (F values: 7.0 – 8.3), midbrain (F values: 8.1 – 8.5), temporal lobe (F values: 7.0 – 8.7) and cerebellar (7.0 – 8.6) activations, activities are also seen to be further extending towards the occipital lobe and visual cortex (F values: 7.0 – 7.4), suggesting potential involvement of visual processing.

These activities spanning from the cortical to deep brain regions are no longer presented in the baseline condition without tFUS stimulation at the thalamus using the same color scale (Fig. 3g-i). The weak electrophysiological activities recorded and shown in the EEG butterfly plot (Fig. 3f) at 83 and 170 ms may also be related to the attention^30^ and/or visual expectation^31^ in the trained monkey.

### tFUS thalamic stimulation remotely elicits V4 activities without visual stimuli

The visual thalamus, particularly the lateral geniculate nucleus (LGN) and pulvinar, plays a crucial role in relaying and modulating visual information to various cortical areas, including area V4. Like we observed the some V4 activations due to tFUS FEF stimulation without presenting actual visual stimuli to the monkey^21^, we also set out to test if we can elicit substantial V4 activities by targeting tFUS stimulation (PRF: 3 kHz, burst duty cycle: 60%) at a region of thalamus (Fig. 1d), i.e., pulvinar. Through the implanted “Utah” array, we were able to assess the tFUS thalamic stimulation effects by remotely recording the neural activities from the V4 area, relying on the substantial connections between the visual thalamus and V4.

After repeating the ultrasound trials randomly in half of all 240 trials, the recorded LFP from one electrode at V4 showed significant differences (*p* < 0.01) during the time window from 22 to 74 ms (Fig. 4a). This demonstrates that without presenting an explicit visual stimulus, the applied steered tFUS (PRF: 3 kHz, DC: 60%) was able to stimulate the visual thalamus and evoke remote neural activity in V4 directly. The inset in Fig. 4a shows a topographical map of t-statistics at 0.1 s, as indicated by the vertical gray dashed line in the LFP waveform. This topomap, covering the 96-channel recording sites over V4, highlights the significant activities (labeled with gray asterisks) elicited by the tFUS stimulation. The purple diamond marks the electrode location selected for the LFP waveforms.

**Figure 4.**
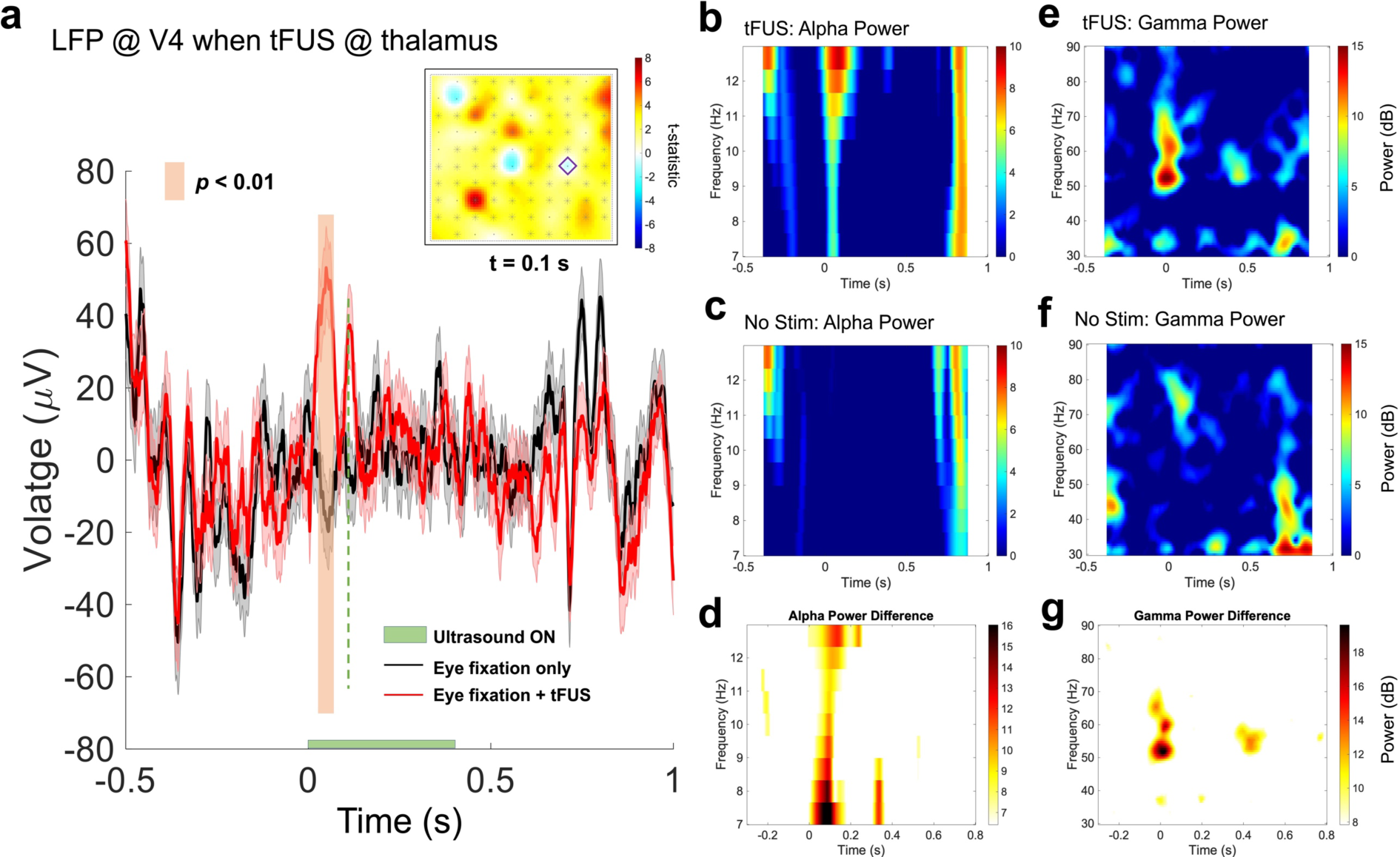
The time-frequency signature of thalamic tFUS-induced V4 activities without visual stimuli. **a**, After repeating 120 trials with ultrasound stimulation (solid red line), a single-channel LFP comparison showed a significantly different time segment versus the eye fixation-only condition (solid black line). The vertical color bar represents the significant difference (*p* < 0.01). The horizontal green bar shows the sonication period, i.e., 400 ms. In the inset, a topographical map of t-statistics at 0.1 s (indicated with a vertical gray dashed line in the LFP waveform) across the 96-channel recording sites over the V4. The purple diamond represents the selected electrode location for the LFP waveforms in (**a**). **b**-**d**, The LFP alpha band (7.5 - 12.5 Hz) contents in the tFUS stimulation condition (**b**), in the eye fixation only condition without stimulation (**c**) and the alpha power difference (**d**). **e**-**g**, The LFP gamma band (30 - 90 Hz) contents in the tFUS stimulation condition (**e**), in the eye fixation only condition without stimulation (**f**) and the gamma power difference (**g**).

It was known that there is pronounced alpha-band synchronization between the pulvinar and V4 in monkeys, which suggested that pulvinar uses the alpha band as a fundamental operating mode to influence cortical areas like V4 during selective attention^33^. We implemented time-frequency analyses across the trials at individual channels, and then took a grand average across the 96 channels. In the comparison of alpha power in the tFUS stimulation (Fig. 4b) versus the no stimulation (Fig. 4c) conditions, there was notable alpha power increase at the beginning of the sonication (9 - 160 ms). The alpha power difference (Fig. 4d) confirmed that by delivering the excitatory tFUS stimulation onto the pulvinar elicits alpha band activities at its connected V4.

It was also suggested that the alpha oscillations contributed to the attention-related effects on gamma frequencies in the V4^33^. We set out to investigate whether there is any time-locked gamma power change at V4 once the tFUS stimulated at pulvinar. Within the gamma band (30 - 90 Hz), when comparing the time-frequency features presented in the tFUS condition (Fig. 4e) vs. those in the trials without ultrasound stimulation (Fig. 4f), the tFUS trials showed significantly increased gamma-band contents averaged across the 96-channel electrode array within 48-71 Hz (Fig. 4g) immediately after the onset of tFUS stimulation, and from 51 to 59 Hz in the time window of 365-504 ms (Fig. 4g, −6 dB contour).

### tFUS thalamic stimulation remotely modulates V4 activities evoked by visual stimuli

Previously, we examined how the excitatory and inhibitory ultrasound stimulation at FEF would remotely modulate the V4 activities evoked by visual stimuli^21^. In this study, we further hypothesized that the tFUS thalamic stimulation effect is also region specific and parameter dependent. For this purpose, we coupled the ultrasound stimulation with the presented visual stimuli, and steered the ultrasound focus along the axial direction to be Δz = +2.4 mm (deeper into pulvinar) or Δz = −2.4 mm (toward the surface of thalamus, e.g., thalamic reticular nucleus, TRN) and tuned the ultrasound pulse repetition frequency to be 3 kHz (excitatory) or 40 Hz (inhibitory).

By applying the 3 kHz PRF tFUS to a spatial location Δz = +2.4 mm at the deep pulvinar, statistically significant modulation (*p* < 0.001) on the visual-evoked LFP waveform was observed within a period of 19 - 303 ms during the sonication and visual stimuli, and 347 – 413 ms near the end of sonication (Fig. 5a). At the time 90 ms post the onset of hybrid stimuli (indicated with a vertical dashed blue line), the significant modulation was extensively presented across the 84 of the total 96 channels in the electrode array (the inset of Fig. 5a) at the implanted location of V4. Given the critical role of alpha-band synchronization between the pulvinar and V4 in monkeys^33^, we are particularly interested in the alpha power changes due to the tFUS neuromodulation. The grand averaged time-frequency representations across the 96 channels obtained from the hybrid stimulation and visual stimulation conditions showed differences mainly at the beginning of sonication, with notably increased spectral power within 10-12.5 Hz (Fig. 5b). When we switched the ultrasound PRF to be 40 Hz at the same location, some significant modulation effects (*p* < 0.001) were maintained, while from the same recording channel, significantly decreased LFP amplitude (*p* < 0.01, Fig. 5c) was also seen after the sonication (650 – 753 ms) was completed. At the same time 90 ms after the stimuli onset, significantly decreased LFP amplitudes were also seen across the recording array (among 49 of the total 96 channels, blue colored area in the inset of Fig. 5c). However, slightly increased alpha power within 8-10 Hz was detected at the beginning of the trial average and after the sonication cessation in the hybrid stimulation condition comparing with that in the visual stimulation condition.

**Figure 5.**
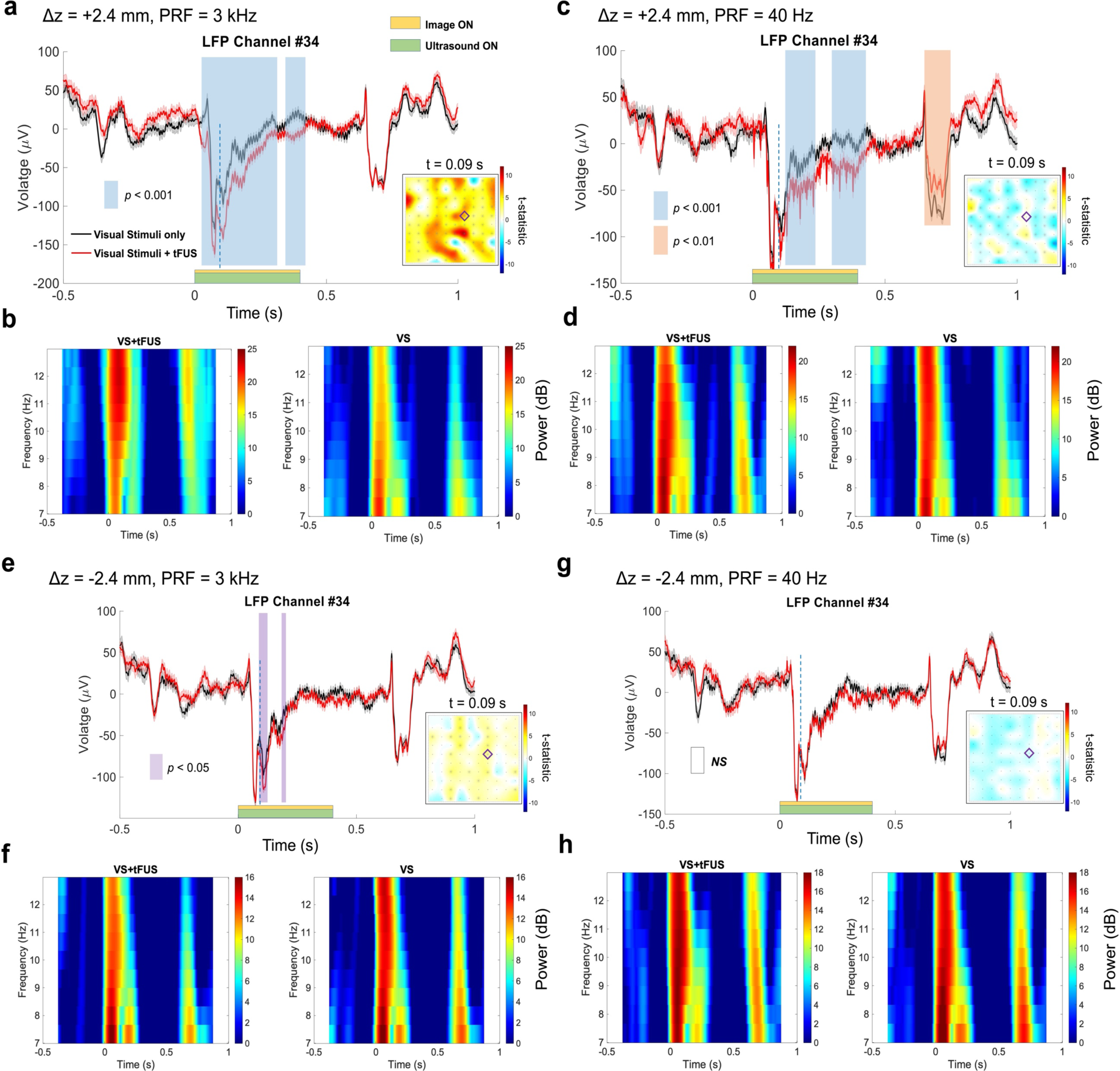
The time-frequency modulation of thalamic tFUS stimulation-induced V4 activities coupled with visual stimuli. **a**-**d**, when tFUS focus was further axially steered by +2.4 mm toward the deep pulvinar from the initial focal location shown in Fig. 1d, the time-frequency modulation by excitatory stimulation using tFUS PRF of 3 kHz (**a**-**b**) or inhibitory stimulation using PRF of 40 Hz (**c**-**d**). After repeating 120 trials with the hybrid stimulation (solid red line), a single-channel LFP comparison showed significantly different time segments versus the visual stimulation only condition (solid black line). The vertical color bar represents the significant difference (blue bars: *p* < 0.001, orange bar: *p* < 0.01). **b** and **d**, The LFP alpha band (7.5-12.5 Hz) contents in the hybrid stimulation condition (left panels) and in the visual stimulation only condition (right panels). **e**-**h**, when tFUS focus was axially steered by −2.4 mm toward the thalamic reticular nucleus (TRN) from the initial focal location shown in Fig. 1d, the time-frequency modulation by excitatory stimulation using tFUS PRF of 3 kHz (**e**-**f**) or inhibitory stimulation using PRF of 40 Hz (**g**-**h**). After repeating 120 trials with the hybrid stimulation (solid red line), a single-channel LFP comparison showed significantly different time segments versus the visual stimulation only condition (solid black line). The vertical color bar represents the significant difference (purple bars: *p* < 0.05). **f** and **h**, The LFP alpha band (7.5-12.5 Hz) contents in the hybrid stimulation condition (left panels) and in the visual stimulation only condition (right panels). The yellow and green horizontal bars in the LFP waveform panels represent the visual and sonication period, i.e., 400 ms, respectively. In the inset, a topographical map of t-statistics at 0.09 s (indicated with a vertical blue dashed line in the LFP waveforms) across the 96-channel recording sites over the V4. The purple diamond in the insets of panel (**a**), (**c**), (**e**), (**g**) represents the same selected electrode location in the “Utah” array as where the LFP temporal waveforms were recorded. In the LFP waveforms, solid colored lines represent the mean, and colored shaded areas demonstrate the S.E.M. Statistics by cluster-based permutation test using Monte-Carlo estimates of the significance probabilities.

By steering the ultrasound focus axially (Δ1z = −2.4 mm) toward the TRN, a shell-like structure that encases the thalamus, and applying the excitatory tFUS stimulation at the PRF of 3 kHz, some statistically significant modulation effects on the visual stimulation evoked LFP waveform within the time windows of 87 – 125 ms and 170 – 205 ms were still observed in the same recording channel, however with a reduced level of statistical significance (*p* < 0.05, Fig. 5e). At t = 90 ms, 20 out of 96 channels exhibited statistically significant difference (the inset in Fig. 5e), which was spatially less extensive than that in the deep pulvinar stimulation. Slightly less averaged alpha power was observed within the frequency window of 10 - 12.5 Hz in the hybrid stimulation condition (left panel in Fig. 5f) than that in the visual stimulation (right panel in Fig. 5f). By switching ultrasound PRF to 40 Hz, no statistically significant change was observed in the LFP waveform (Fig. 5g), across the electrode array (the inset in Fig. 5g), and in the spectral features (Fig. 5h).

## Discussion

In the present study, we showed that tFUS can be used to target and modulate the visual thalamus and to investigate the resultant effects on interconnected cortical and deep brain areas involved in the visual and cognitive processing through multi-channel EEG source imaging of whole brain and intracranial electrophysiological recordings specifically at V4. Through the *in vivo* experiments on the behaving monkey model, we were able to assess the brain stimulation effects by the tFUS stimulation only or by the hybrid stimulation of ultrasound coupled with visual stimuli. By targeting the ultrasound stimulation at the thalamus, we also observed that the induced neural effects on the visual-evoked local field potential at V4 are also region specific and parameter dependent similar to what we previously noted in the FEF stimulation^21^.

In this study, we harnessed the ESI to localize the brain activities that were time-locked to the ultrasound stimulation events, which provided valuable insights of the spatial locations of tFUS stimulation evoked brain responses. We observed that the FEF or thalamic tFUS stimulation elicited initial brain responses localized at the respective targeted area (Figs. 2 and 3). We adjusted the ESI reconstruction method from minimum norm for cortical source estimation to sLORETA for deep brain source imaging in our data analysis pipeline. We had a pilot study testing the feasibility of using scalp EEG source imaging method to monitor the brain responses to low-intensity tFUS stimulation in an anesthetized rat model^34^. As a comparison, the present work further tested the feasibility of ESI in a head-fixed behaving large animal model which provided a noninvasive brain imaging opportunity to examine the cortical and deep brain responses to the tFUS stimulation in a more natural setting than in the functional magnetic resonance imaging. Further investigations are warranted for connectivity analyses based on high-density scalp EEG source imaging^23,35,36^ results to better understand the potential local and global changes^19^ due to the tFUS stimulation at the visual thalamus.

We took the advantages of implanted “Utah” array recording remotely from the site of ultrasound stimulation, i.e., FEF or thalamus, and eliminated numerous confounding factors commonly associated with physical electrode-based electrophysiological recordings, such as potential electrode vibration^37^. By avoiding artifacts in the recordings and neural effects induced by electrode vibrations, this is deemed as a rigorous approach to study tFUS-induced effects in a well-defined visual brain network without the interference of mechanical vibrations. However, to study the cell-type specificity in the visual thalamus, like the pulvinar, by directly recording from this deep brain region would shed light on identifying effective ultrasound parameters, e.g., PRF^10^ and duty cycle^38^, to induce more explicit excitatory or inhibitory effects in the visual processing. The existence of excitatory neuron types, e.g., relay cells projecting to the visual cortical areas, such as V4 and other higher-order cortical regions and inhibitory GABAergic interneurons within the pulvinar^39^, may provide an important testing bed for further delineating the functional cell-type responses to the tFUS stimulation at the visual thalamus. It would also be of tremendous interests to test how the reticular thalamic neurons and corticothalamic neurons would be modulated by the tFUS parameters in terms of sensory gating and the thalamic modulation in response to attentional demands and optimizing the stimulation paradigm to enhance the attentional processes. To address these unmet needs, acoustically transparent electrodes enabled with flexible electronics might be highly helpful to have implantations directly into the thalamus and its interconnected brain regions, such as superior colliculus, to measure and assess the impact of thalamic tFUS stimulation on the visual attention and sensory processing.

The beam steering capability of the customized 128-element random array ultrasound transducer had enabled us to scan through the depth of FEF and test the subregional specific neuromodulation effects on the visual-evoked potentials at V4^21^. In the present study, we further employed this technological feature to axially change the spatial location of ultrasound focus within the thalamus and evaluate the change of neuromodulation effects. The results in Fig. 5 demonstrated the regional specific effects and validated the substantial thalamocortical connections between the pulvinar and V4.

As a negative control, we delivered tFUS stimulation to the insula^21^ by positioning the random array ultrasound transducer at the same scalp location with different axial steering from the thalamic stimulation. The insula tFUS stimulation did not generate any significant modulation in the LFP waveforms at V4, which confirmed the dependence of remote neural effects on the original stimulation site. This control study also ruled out the potential auditory side effects^40,41^, induced by the PRF^42^.

By targeting the thalamus using noninvasive tFUS stimulation, we intend to contribute valuable insights into the neural underpinnings of visual processing and the therapeutic potentials of ultrasound-based neuromodulation for treating visual, sensory and cognitive impairments. To evaluate the safety of thalamic tFUS stimulation, we monitored the temperature at the sonication site using a circuit board (AD8495, Adafruit Industries, New York, NY, USA) connected to a thermocouple placed on the scalp during the stimulation. The observed temperature increase was less than 0.01°C. We also assessed the subject’s behavior for ten days before and ten days after the tFUS sessions. After repeated tFUS sessions, the subject showed no signs of adverse effects. The average water intake during task paradigms before and after thalamic tFUS stimulation did not show significant changes, which indicates that the monkey’s behavioral state remained stable.

Overall, our study demonstrated that the thalamic tFUS can produce substantial neuromodulatory effects in cortical and deep brain regions evidenced through multiscale electrophysiological recordings. Moving forward, it is essential to conduct a more in-depth exploration of the intricate behavioral changes resulting from such a noninvasive thalamic stimulation by finely tuning pressure and PRF levels. This includes examining the path and speed of eye movements, as well as the nuances of visual perception, that arise from administering tFUS with more extensive parameter configurations and targeted subregions. By analyzing behavior in conjunction with neural recordings, we can gain a more profound understanding of the far-reaching neuromodulatory effects of tFUS, ultimately propelling its advancement as a potent neuromodulation technology.

## Data availability

The data are presented in the paper. Additional data will be made public through a data repository upon paper acceptance.

## Acknowledgments

This work was supported in part by NIH grants RF1NS131069, U18EB029354, and R01NS124564. We would like to thank Dr. Matthew Smith for supporting the data collection in nonhuman primates in his laboratory, and Samantha Schmitt, Yunruo Ni and Emily Crane for their work in collecting the data in the Smith Laboratory and assisting with post-processing. We would also like to thank Dr. Zhengxiang Cai for useful discussion on source imaging. Cartoons in Figure 1a were created with BioRender.com.

## Competing Interest Declaration

B.H. and K.Y. are co-inventors of a pending patent application for tFUS technique.

## Notes

### Competing Interest Statement

The authors are co-inventors of a pending patent application for tFUS technique.

